# Validation of an AI-Powered Automated Colony Analysis Platform Across Eight ISO Microbiological Methods: A Multi-Pathogen, Multi-Matrix Performance Study

**DOI:** 10.64898/2026.05.08.723721

**Authors:** Jennifer Upfold, Aimee van de Schoor, Helena Falholt Elvebakken, Oscar Petersen, Christopher Falholt, Carina Küstner, Magnus Madsen

## Abstract

Manual colony counting remains the rate-limiting, operator-dependent step in culture-based food microbiology quality control (QC). Automated colony analysis using machine learning (ML) offers the potential to standardise, accelerate, and improve the traceability of this process. However, systematic multi-method validation data for AI-based platforms against recognised international standards remain scarce. We conducted a prospective, multi-study validation of the Reshape Smart Incubator which is an automated imaging and ML-based colony analysis system, across eight ISO microbiological reference methods. In total, 887 plates were analysed, spanning qualitative (presence/absence) detection of *Listeria* spp. (ISO 11290-1) and *Salmonella* spp. (ISO 6579), and quantitative enumeration of total viable count (ISO 4833), *Bacillus cereus* (ISO 7932), *Enterobacteriaceae* (ISO 21528), coagulase-positive *Staphylococci* (ISO 6888), yeasts and moulds (ISO 21527), and lactic acid bacteria (ISO 15214). Automated results were benchmarked against the consensus of three or more trained technicians. The platform achieved 100% agreement with manual assessment for all both qualitative detection methods (ISO 11290-1, ISO 6579) with zero false positives and zero false negatives. For quantitative enumeration, agreement ranged from 92.97% (ISO 15214, n=122, using ISO-aligned ±10%/>30 CFU thresholds) to 98.46% (ISO 21528, n=130). Where discrepancies occurred, they largely coincided with plates showing high inter-technician variability. Precision testing demonstrated a coefficient of variation of 5.88% and a mean standard deviation of 0.44 CFU for low-count plates. This study presents a comprehensive multi-ISO validation of an AI-based colony analysis system to date. The AI models demonstrated performance comparable to or exceeding that of trained human technicians across a broad range of microbiological targets, agar types, and colony morphologies, thereby supporting their use as a validated and traceable alternative to manual plate reading in accredited food microbiology quality control laboratories.

## 1. Introduction

Culture-based microbiological testing remains the cornerstone of food safety surveillance and quality control worldwide (1,2). From raw ingredient testing to finished product release, regulatory frameworks in the European Union, United States, and internationally mandate the enumeration and detection of specific indicator organisms and pathogens using internationally harmonised reference methods, principally those published by the International Organization for Standardization (ISO).

Central to these workflows is the manual reading of agar plates, during which a trained technician inspects incubated Petri dishes, counts individual colonies, identifies morphologies relevant to the target organism, and determines whether a plate should be classified as positive, negative, or too numerous to count (TNTC) (1). This process is labour-intensive, time-consuming, and inherently subjective. Studies have consistently documented significant inter-operator variability in colony counts, particularly for small or overlapping colonies, morphologically ambiguous species, and plates near the limits of countable ranges (2,3). Such variability introduces uncertainty into batch release decisions, complicates inter-laboratory comparisons, and can undermine data integrity in regulated environments (2,4).

Automated colony counting systems have existed in various forms for several decades, ranging from early threshold-based image analysis systems to more recent machine learning-based platforms (6-12). However, adoption in accredited quality control laboratories has remained limited due to the lack of robust validation data demonstrating equivalence to manual methods across the breadth of ISO-defined workflows (10,11). Most published validation studies have focused on single organisms, specific agar types, or narrow laboratory applications, with few studies assessing performance across both qualitative pathogen detection and quantitative enumeration simultaneously.

The Reshape Smart Incubator (Reshape Biotech, Copenhagen, Denmark) is an integrated platform combining automated plate imaging under controlled and reproducible lighting conditions with proprietary machine learning models for colony detection, classification, and enumeration. The system generates structured, auditable outputs with full imaging traceability, features directly relevant to laboratory accreditation under ISO/IEC 17025.

The aims of this study were therefore threefold: (1) to evaluate prospectively the Reshape Smart Incubator against eight internationally recognised ISO microbiological reference methods spanning both qualitative pathogen detection and quantitative enumeration across organism groups relevant to food safety quality control; (2) to characterise the relationship between inter-analyst variability and automated system discrepancy in order to distinguish genuine model error from intrinsic plate ambiguity; and (3) to assess the repeatability and cross-device reproducibility of the automated platform under conditions representative of routine laboratory use. Collectively, these aims address key evidence gaps that have thus far limited the adoption of automated imaging systems in accredited food microbiology laboratories.

## 2. Materials and Methods

### 2.1 Study Design and Overview

Eight independent, prospective validation studies were conducted, each targeting a specific ISO method. All studies shared a common structural design: (i) preparation of known positive and/or negative strains across a range of dilutions spanning both the countable range and the uncountable range (too numerous to count; TNTC), to ensure the model was trained to recognise and correctly classify plates at both extremes of colony density; (ii) plating onto the selective or non-selective agar specified by the respective ISO method; (iii) incubation under ISO-specified conditions; (iv) automated imaging using the Reshape Smart Incubator; and (v) independent manual counting by a minimum of three trained personnel, blinded to the automated result. Final performance metrics were calculated against the consensus of trained assessors.

Studies were designed to simulate the range of plate types and colony densities encountered in routine QC workflows, including both pure cultures and mixed-species plates where specified by the method. Automated plate analysis was performed using the Reshape Smart Incubator without modification of the underlying microbiological method at any stage.

### 2.2 Automated Imaging System

The same plates assessed manually by trained personnel were imaged using the Reshape Smart Incubator (Reshape Biotech, Copenhagen, Denmark), an automated incubation and imaging platform designed for standard microbiological workflows. The system captures high-resolution plate images under controlled and reproducible epi-illumination (top light) and transillumination (bottom light) conditions to minimise variability arising from ambient lighting and plate positioning.

Images were analysed using convolutional neural network (CNN)- and transformer-based models trained for colony detection, classification, and enumeration across selective and non-selective microbiological media. Colony detection was formulated as an object detection and counting task, with model outputs aggregated at the plate level. Depending on the ISO method, models were configured either as general-purpose enumeration models or as method-specific classifiers optimized for characteristic colony morphologies and chromogenic responses associated with the target organism. Model outputs included total colony count, colony localisation, TNTC (too numerous to count) classification, and where applicable, binary positive/negative classification for qualitative methods. All images and associated inference outputs were stored with version-controlled audit trails to support traceability and reviewability in regulated laboratory environments.

For ISO 4833 (Total Viable Count), analysis was performed using Microbiology Model v2.4.1, a transformer based general-purpose enumeration model trained across multiple non-selective and semi-selective agar types. For all other methods, method-specific models trained on the corresponding selective or chromogenic media were used. Full model version information is provided in Supplementary Table S1.

### 2.4 Model Training and Validation Independence

All models were trained prior to validation using independent development datasets distinct from the validation datasets reported here. Validation was performed prospectively using fixed model weights without post hoc retraining or manual adjustment. Manual assessors were blinded to automated results throughout the study. Training data included diverse historical plate images spanning multiple agar types, colony morphologies, colony densities, and imaging conditions, with standard augmentation procedures applied to improve robustness.

### 2.5 Reference Methods and Organisms

A summary of the the eight ISO methods evaluated, the target organisms, method type, total number of plates analysed, and key performance results are presented in Table 1. Full strain details, culture conditions, and agar preparation protocols for each method are provided in Supplementary Table S1.

**Table 1.** Summary of ISO method validation studies conducted on the Reshape Smart Incubator platform.

| ISO Method | Target Organism | Method Type | n (plates) | Agreement (%) | Key Metrics |
| --- | --- | --- | --- | --- | --- |
| ISO 11290-1 | Listeria spp. | Qualitative | 120 | 100% | 0% FP, 0% FN; 70.83% TP, 29.17% TN |
| ISO 6579 | Salmonella spp. | Qualitative | 85 | 100% | 0% FP, 0% FN; 28% TP, 72% TN |
| ISO 6888 | Coagulase-pos. Staphylococci | Quantitative | 87 | 100% | 0% FP, 0% FN; 11% TP, 89% TN |
| ISO 4833 | Total Viable Count | Quantitative | 157 | 98.5% | CV=5.88%; SD=0.44 CFU (low count) |
| ISO 7932 | Bacillus cereus | Quantitative | 114 | 98.25% | 14 TNTC TP, 100 Not-TNTC TN; 0 FP/FN |
| ISO 21528 | Enterobacteriaceae | Quantitative | 130 | 98.46% | 0% FP, 1% FN (1 miscount) |
| ISO 21527 | Yeasts & Moulds | Quantitative | 138 / 119 | 92.75% / 97.48% | Yeast / Mould |
| ISO 15214 | Lactic acid bacteria | Quantitative | 122 | 92.97% | FP=0%; FN=3% (6 TNTC miscl.) |

FP = false positive; FN = false negative; TP = true positive; TN = true negative; CV = coefficient of variation for plate orientation; SD = standard deviation; TNTC = too numerous to count.

### 2.6 Qualitative Methods (ISO 11290-1 and ISO 6579)

*ISO 11290-1:2017 (Listeria spp*.*): Listeria monocytogenes* was inoculated into brain heart infusion (BHI) broth and grown overnight at 37°C. After dilution to the countable range, strains were plated alone and in combination with other microorganisms to simulate background contamination. Plating was performed on chromogenic Listeria agar (CLA) and plates were incubated at 37°C for 24–48 hours. Presumptive *Listeria* colonies on CLA appear as characteristic blue-green colonies, with or without an opaque halo. In total, 120 plates (or images of plates) from seven independent assays were evaluated using three independent manual assessors.

*ISO 6579-1:2017 (Salmonella spp*.): was inoculated into BHI broth and grown overnight at 37°C. Suspected-positive and confirmed-positive enrichment broths were plated onto Xylose-Lysine-Deoxycholate (XLD) agar and incubated at 37°C for 24 hours. Presumptive *Salmonella* colonies appear red with or without a black centre on XLD. Plates were prepared as pure cultures and in combination with [background organism] to assess model specificity under mixed-flora conditions. A total of 85 plates were evaluated using three independent manual assessors.

For both qualitative methods, classification was binary (positive/negative) and concordance was assessed against the manual consensus. Confusion matrices (true positive, true negative, false positive, false negative) were calculated at the plate level.

### 2.7 Quantitative Methods (ISO 4833, 6888, 7932, 15214, 21527, 21528)

For all quantitative methods, plates were prepared from decimal dilution series of pure or mixed cultures at concentrations designed to yield plates within the ISO-defined countable range (typically 15 to 300 CFU per plate) and above the upper countable limit (TNTC). Automated colony counts were compared to the mean of three or more independent manual assessors. Agreement was defined per ISO-aligned thresholds: ±10% for plates above 30 CFU, and ±3 CFU for plates with 30 or fewer colonies. TNTC classification accuracy was assessed via confusion matrix for all quantitative methods, with the TNTC threshold defined per the relevant ISO standard (300 CFU for ISO 4833 and ISO 21528; 150 CFU for ISO 6888).

Specific agar media used: Baird-Parker agar (ISO 6888); plate count agar (ISO 4833); Mannitol Egg Yolk Polymyxin agar (ISO 7932); MRS agar (ISO 15214); DRBC or DG18 agar (ISO 21527); VRBG agar (ISO 21528).

### 2.8. Inter-Analyst Variation Analysis

The dataset consisted of colony count data generated using the ISO 4833 and ISO 15214 quantitative microbiological methods, both of which involve challenging manual enumeration of diverse colony morphologies, including bacteria, yeast, and fungi distributed both on the agar surface and within the media. Inter-analyst variability for each plate was quantified as the standard deviation of all available human counts, while human consensus was defined as the median analyst count. Model deviation was calculated as the absolute difference between the ML prediction and the human consensus. Plates that were TNTC were excluded from analysis.

### 2.9. Repeatability and Cross-Device Reproducibility

Precision of the automated system was assessed for ISO 4833 by re-imaging 15 representative plates 12 times each under varied plate orientations, translations, and across multiple devices. Repeatability was quantified as the standard deviation (for plates with 30 CFU or fewer) and coefficient of variation (for plates with more than 30 CFU).

### 2.10 Statistical Analysis

For qualitative methods, primary performance metrics were sensitivity, specificity, and overall concordance with manual assessment. For quantitative methods, the primary metric was the percentage of plates falling within ISO-aligned agreement thresholds. Secondary metrics included Pearson correlation coefficient (r) between automated and manual counts, log_10_(CFU/mL) correlation, and TNTC classification accuracy. Associations between inter-analyst variability and model deviation were assessed using Spearman’s rank correlation, and plates were additionally stratified by disagreement level for comparison using a Kruskal-Wallis rank-sum test.

## 3. Results

### 3.1. Qualitative Detection: *Listeria spp*. and *Salmonella spp*

The platform achieved 100% concordance with manual assessment for both qualitative detection methods (Table 2). For ISO 6579 (*Salmonella* spp., n=85 images), zero false positives and zero false negatives were recorded across seven independent assays involving three separate human assessors. True positive and true negative rates were 28% and 72% respectively. The higher proportion of true negatives in this dataset reflects the design of the study, which included a deliberate enrichment of negative controls to challenge the model’s specificity. For ISO 11290-1 (*Listeria* spp., n=120 images), the model again achieved 100% concordance. True positives accounted for 70.83% and true negatives for 29.17% of classified plates.

**Table 2.** Confusion matrix summary for qualitative (presence/absence) detection methods.

| Organism | n (plates) | True Positive | True Negative | FP/FN |
| --- | --- | --- | --- | --- |
| Salmonella spp.<br>(ISO 6579) | 85 | 28% | 72% | 0% / 0% |
| Listeria spp.<br>(ISO 11290-1)) | 120 | 70.83% | 29.17% | 0% / 0% |

*FP* = false positive; *FN* = false negative; *TP* = true positive; *TN* = true negative.

### 3.2. Quantitative Enumeration: coagulase-positive *Staphylococci* (ISO 6888)

Evaluation against ISO 6888 (n=87 plates, Baird-Parker agar) demonstrated 100% agreement between automated and manual counts. TNTC classification was perfect: all 11% of TNTC plates were correctly identified by the model and all 89% of countable plates were correctly classified as not-TNTC (FP=0%, FN=0%). Colony counts across the countable range showed tight clustering around the line of perfect agreement.

### 3.3. Quantitative Enumeration: ISO 4833 (Total Viable Count)

The ISO 4833 dataset (n=112 plates) included diverse colony morphologies (bacterial, yeast, mould), surface and pour-plate inoculation methods, and real-world artefacts including plate labels and barcodes. Applying ISO-aligned agreement thresholds (plus or minus 10% for plates above 30 CFU; plus or minus 3 CFU for plates at or below 30 CFU), the automated system achieved 98.5% agreement.

### 3.4. Quantitative Enumeration: ISO 7932 (Bacillus cereus)

Validation against ISO 7932 (n=114 plates, MYP agar) achieved 98.25% agreement. The study included both *B. cereus* (NCIMB 9373, pink-orange colonies with lecithinase halo) and *B. subtilis* (NCIMB 13061, yellow colonies without halo), challenging the model to differentiate morphologically distinct colony types on the same selective medium. TNTC classification was perfect: 14 TNTC plates were all correctly identified, and all 100 non-TNTC plates were correctly classified (FP=0%, FN=0%). Analysis indicated that agreement was higher under top-light (epi-illumination) versus bottom-light (transillumination) conditions, a finding that has been incorporated into updated operating recommendations.

### 3.5. Quantitative Enumeration: ISO 21528 (Enterobacteriacea)

The ISO 21528 study (n=130 plates, VRBG agar) used three species: *Serratia marcescens, Escherichia coli* NCIMB 12805, and *Bacillus subtilis* NCIMB 13061, including mixed-species plates to simulate real-world sample complexity. Overall agreement was 98.46%. The confusion matrix showed 0% false positives, 1% false negatives (corresponding to one misclassified plate), 32% true positives (TNTC), and 67% true negatives. In cases of high inter-rater variability among human counters, the automated count was more consistent than the human mean, suggesting the system provides a stabilising effect on QC data quality.

### 3.6. Quantitative Enumeration: ISO 21527 (Yeast and Moulds)

Yeasts and moulds present a unique challenge for automated counting because fungal colonies can spread across plates, overlap, and are substantially more heterogeneous in morphology than bacterial colonies. For ISO 21527 (DRBC or DG18 agar, 39 assays, n=138 yeast images and 119 mould images), the platform achieved 92.75% agreement for yeast enumeration and 97.48% for mould enumeration. The higher performance for moulds likely reflects the more visually distinct and spatially constrained colony morphologies on DRBC agar. These results demonstrate that deep learning-based approaches can meaningfully overcome the inter-observer variability and counting errors that have historically plagued manual yeast and mould enumeration.

### 3.7. Quantitative Enumeration: ISO 15214 (Lactic Acid Bacteria)

Evaluation against ISO 15214 (n=122 images, MRS agar, three LAB species: *Levilactobacillus brevis, Pediococcus pentosaceus*, and *Lactiplantibacillus plantarum*) yielded 92.97% overall agreement. The confusion matrix showed 0% false positives, 3% false negatives (six plates classified as countable by the model but as TNTC by trained personnel), 38% true positives, and 59% true negatives. The small number of TNTC misclassifications likely reflects the high colony density and colony clustering characteristic of LAB on MRS agar at the TNTC boundary, and is expected to improve with additional training data.

### 3.8. Inter-Analyst Variation Analysis

A strong positive association was observed between inter-analyst variability and model deviation from human consensus (Figure 1; Spearman’s ρ = 0.695, p < 2.2×10^−16^). Plates associated with higher analyst disagreement exhibited substantially greater model error, suggesting that model deviations were concentrated on inherently ambiguous or difficult samples rather than arising from systematic independent model bias.

**Figure 1.**
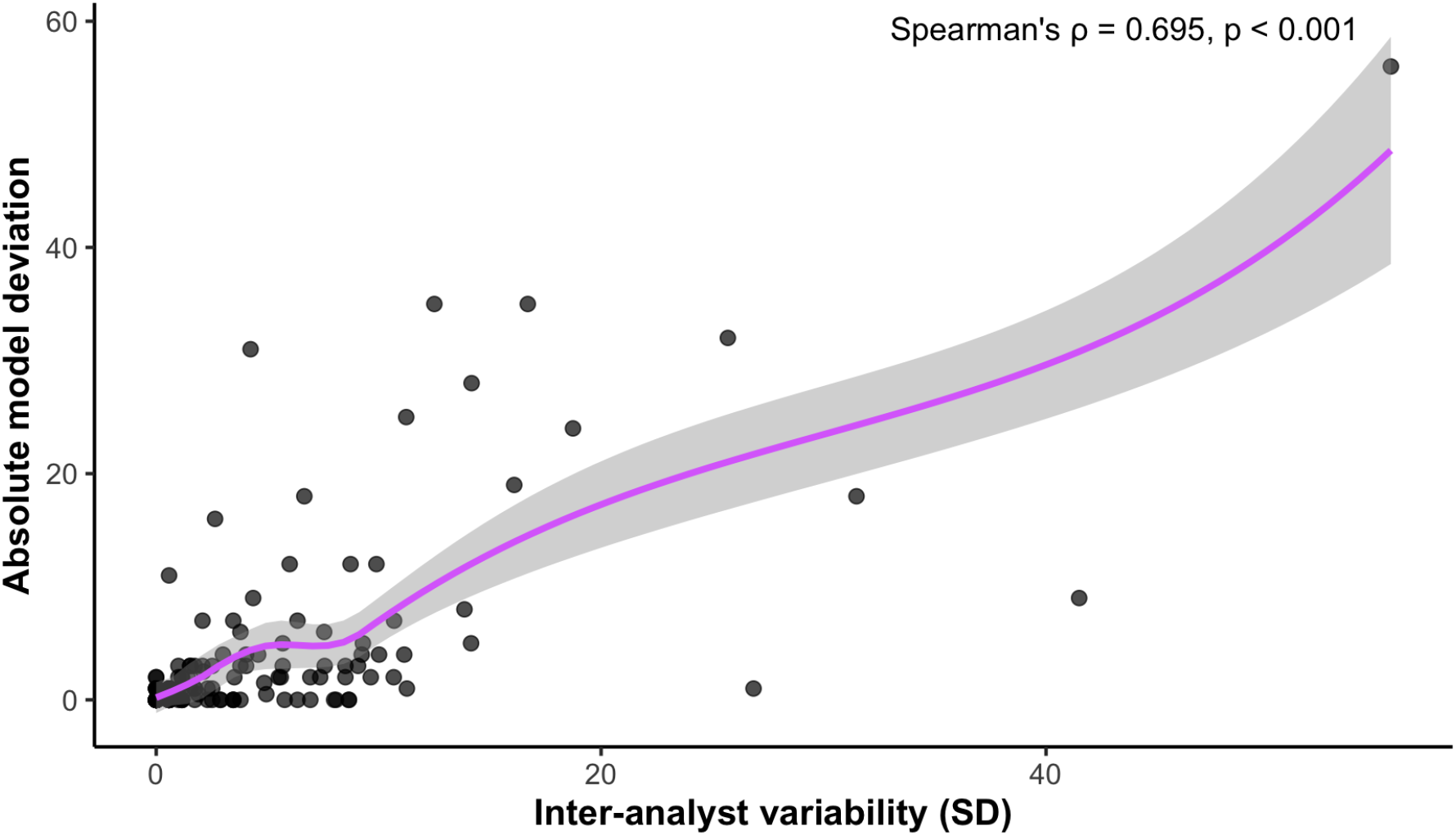
Association between inter-analyst variability and ML model deviation across ISO 4833 and ISO 15214 plate counts. Model deviation increased with inter-analyst variability, indicating that larger model errors occurred primarily on plates where human analysts also disagreed, suggesting that reduced performance was mainly driven by plate complexity rather than model instability.

### 3.9. Repeatability and Cross-Device Reproducibility

A subset of 15 plates (spanning the full countable range) from the ISO 4833 dataset was selected to assess the repeatability and cross-device reproducibility. Each plate was re-imaged 12 times under systematically varied conditions including plate orientation, translational position within the imaging chamber, and across multiple devices.

For low-count plates (30 CFU or fewer), the mean standard deviation across replicate imaging sessions was 0.44 CFU, indicating that the system produces essentially identical counts regardless of how the plate is positioned or which device is used. For higher-count plates (above 30 CFU), the mean coefficient of variation across replicate sessions was 5.88%, confirming stable proportional performance across the countable range.

## 4. Discussion

### 4.1 Automated Colony Enumeration Achieved Human-Level Performance Across Diverse ISO Methods

Across 887 plate images and eight ISO reference methods, the Reshape ML models demonstrated performance at or above the level of trained human technicians. For qualitative detection, which is one of the highest-stakes applications of microbiological plate reading, where a false negative can lead to the release of contaminated product, the platform achieved 100% concordance with zero false negatives and zero false positives across three independent methods and organisms. This is arguably the most important finding of this study: an AI system can be trusted to make binary safety-critical decisions at least as reliably as a trained microbiologist.

An additional notable finding was the platform’s strong performance at low colony concentrations, where automated image-analysis systems have historically been prone to false-positive detections caused by artefacts such as air bubbles, agar imperfections, condensation, reflections, or dish imprints (5-9,12,13). Across both Salmonella and Listeria qualitative workflows, the Reshape ML models maintained complete specificity even at low target concentrations, demonstrating that the underlying vision models were able to distinguish true microbial growth from non-biological imaging artefacts. This is particularly relevant for food safety applications, where false-positive classifications can trigger unnecessary investigations, product holds, and confirmatory testing

For quantitative enumeration, agreement rates ranged from 92.97% to 98.5% depending on the method. The lower end of this range (ISO 15214, Lactic Acid Bacteria) reflects both the methodological challenge of counting diverse colony morphologies across a wide concentration range, especially given the diffuse or irregular colonies that can be poorly separated in the agar for this method, as well as the rigorous ISO-aligned thresholds applied. When agreement is assessed relative to individual human counters rather than the consensus, the automated count frequently falls within the range of at least one assessor, suggesting that remaining discrepancies represent inherent biological ambiguity rather than model failure. This mirrors the experience of laboratory proficiency testing programmes, in which colony count discrepancies between experienced analysts on the same plate are routinely observed (3,14).

Previous generations of automated colony counters relied primarily on threshold-based segmentation or morphology filtering and were often limited by poor performance on crowded plates, overlapping colonies, or heterogeneous morphologies (5,6,12,13). More recent machine learning-based approaches have improved detection capability but have generally been evaluated using narrow datasets or single-organism workflows (7,8,10,11). In contrast, the present study evaluated performance prospectively across eight ISO reference methods spanning both qualitative pathogen detection and quantitative enumeration, thereby providing a broader assessment of operational suitability for accredited food microbiology laboratories.

### 4.2 Model Deviations Were Associated with Intrinsically Ambiguous Plates

The strong positive association observed between inter-analyst variability and machine learning model deviation indicates that larger automated discrepancies occurred primarily on plates that were also difficult for human analysts to interpret. Given the complexity of methods such as ISO 4833 and ISO 15214, these findings suggest that reduced model agreement was driven predominantly by intrinsic plate ambiguity rather than systematic model instability.

The automated system showed minimal deviation on plates where human analysts strongly agreed, whereas higher disagreement between analysts was associated with substantially greater model deviation. This finding has direct operational relevance for quality control laboratories. The plates generating the greatest uncertainty for the automated system are the same plates that would typically prompt recounting or supervisory review during conventional manual analysis. As a result, expert attention can be directed precisely toward the most ambiguous cases while maintaining consistency across routine high-throughput workflows.

### 4.3 Automated Enumeration Demonstrated High Repeatability and Cross-Device Reproducibility

To evaluate repeatability and cross-device reproducibility, 15 plates spanning the full countable range of the ISO 4833 dataset were re-imaged 12 times under varied imaging conditions, including changes in orientation, positioning, and device. For plates with ≤30 CFU, the mean standard deviation across replicate imaging sessions was 0.44 CFU, demonstrating highly consistent counts independent of plate placement or device. For plates with >30 CFU, the mean coefficient of variation was 5.88%, confirming stable proportional performance across the countable range.

These results demonstrate that the automated system’s output is independent of operator handling, plate positioning, and device-to-device variation. This degree of within-system consistency is a prerequisite for deployment in accredited laboratories operating under ISO/IEC 17025, where measurement reproducibility must be demonstrable and documented, and is difficult to achieve through manual counting where individual handling differences inevitably introduce variation.

### 4.3 Operational and Traceability Advantages

Beyond analytical agreement, automated colony analysis offers several operational advantages that are not fully captured by performance statistics alone. First, the system generates high-resolution, version-controlled images of every analysed plate, creating a fully auditable record of the raw data underlying each quality control decision. This substantially strengthens traceability and reviewability compared with conventional manual counting workflows, where the physical plate itself is often the only retained record.

Second, imaging and analysis are decoupled from the specific operator performing the assessment, reducing reliance on experienced personnel for routine plate reading and enabling expert microbiologists to focus on higher-value tasks such as investigations, troubleshooting, and method optimisation. Third, the software generates structured digital outputs that can integrate directly with laboratory information management systems (LIMS), thereby reducing transcription errors and accelerating result reporting.

### 4.4 Limitations and Future Work

Several limitations should be considered when interpreting the present findings. First, validation was performed under controlled laboratory conditions using known strains at defined concentrations. Although this approach enables rigorous benchmarking, additional external validation is required to assess performance under routine operational conditions. Ongoing studies are therefore evaluating the platform across multiple external laboratories using predefined pass/fail success criteria designed to assess robustness, reproducibility, and usability outside of a controlled in-house laboratory environment.

Second, although the datasets were sufficient to evaluate overall performance trends, larger datasets would permit more precise characterisation of agreement across the full count distribution, particularly at the extreme upper and lower limits of quantification.

Future work will focus on extending method coverage to ISO 11290-2 (Listeria enumeration), and integrating the platform with automated sample preparation systems to enable streamlined workflows from dilution through final result generation.

## 5. Conclusions

This study presents the first systematic multi-ISO validation of an AI-based colony analysis platform across a broad range of culture-based food microbiology quality control workflows. The Reshape platform (Smart Incubator and vision AI models) achieved 100% concordance with expert manual assessment for all qualitative detection methods, including the major foodborne pathogens *Listeria monocytogenes* and *Salmonella* spp., while also demonstrating consistently high and reproducible performance across six quantitative enumeration methods.

Taken together, these findings provide a strong scientific basis for the adoption of automated colony imaging as a validated and traceable alternative to manual plate reading in accredited food microbiology laboratories. The combination of expert-level agreement, high reproducibility, full imaging traceability, and reduced operator-dependent variability positions automated colony imaging as a promising advancement for food microbiology quality control.

## Data Availability

Raw plate image data and colony count datasets supporting the findings of this study are available from the corresponding author upon reasonable request.

