## Supplemental Figure S1 & S2 for "Validation of an AI-Powered Automated Colony Analysis Platform Across Eight ISO Microbiological Methods: A Multi-Pathogen, Multi-Matrix Performance Study"

**Supplementary Figures**
Supplementary figures for validation statistics presented in the manuscript.

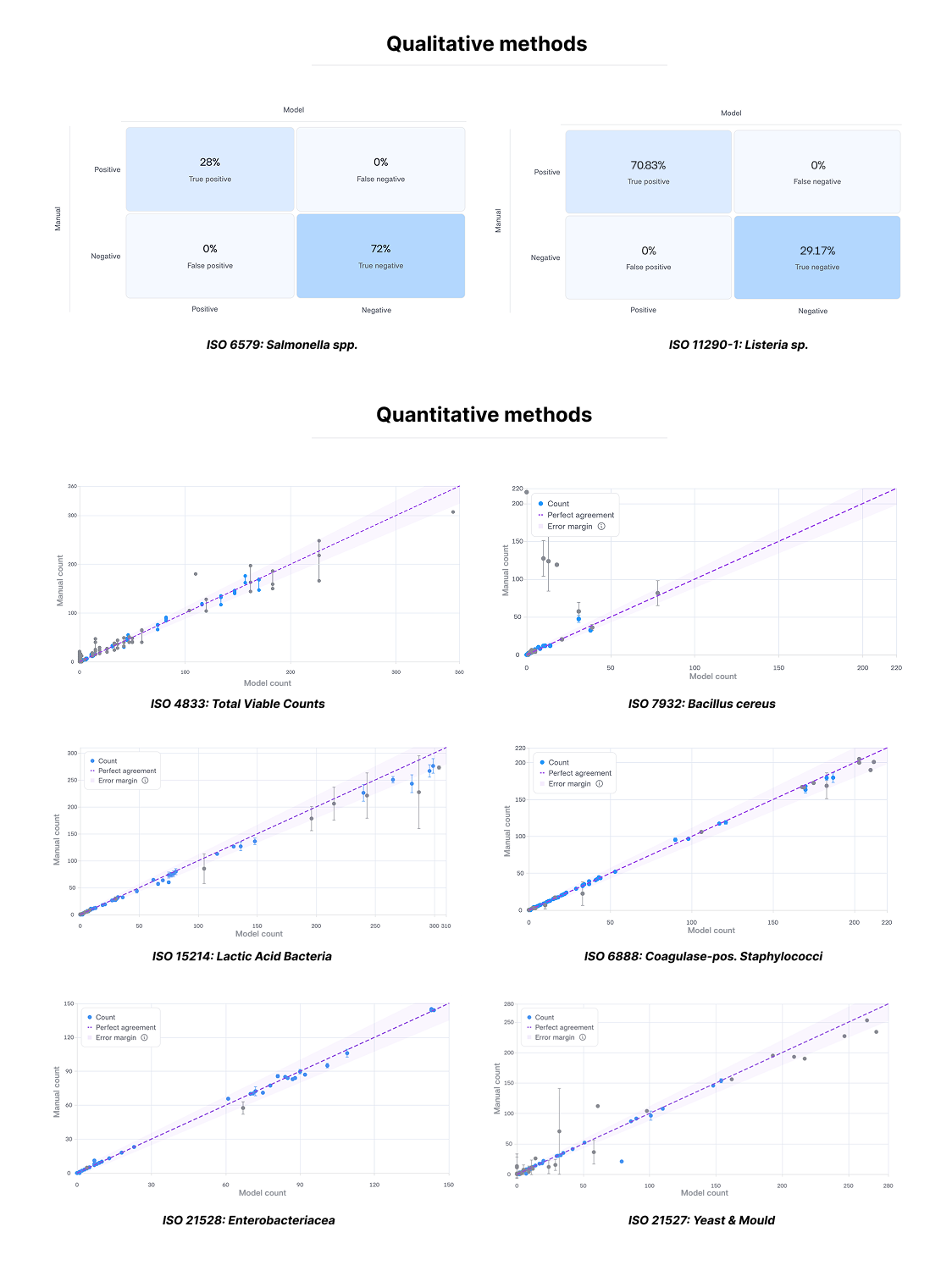

**Figure S1.** Automated system performance across two qualitative ISO reference methods, presence/absence detection (ISO 6579, *Salmonella* spp.; ISO 11290-1, *Listeria* sp.).

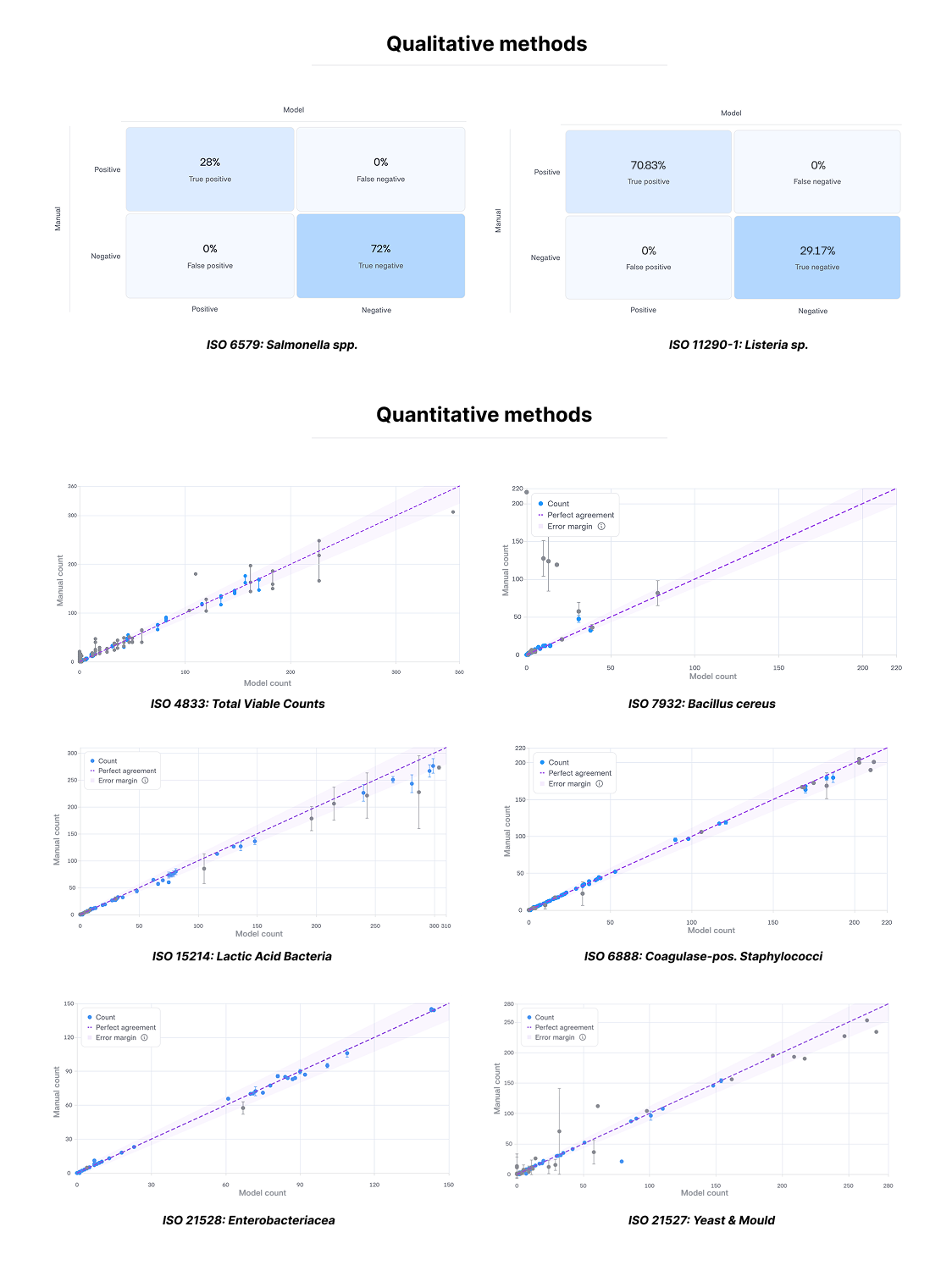

**Figure S2.** Automated system performance across six ISO reference methods. Correlation plots of model counts versus manual consensus counts for six quantitative enumeration methods. Dashed purple line indicates perfect agreement; shaded area indicates the ISO-aligned error margin. Grey points indicate plates excluded from statistics due to high inter-analyst variability.
